# hnRNPA2B1 Modulates Early TIA1 Recruitment into Stress Granules through an RNA-Dependent Assembly Mechanism

**DOI:** 10.64898/2026.04.22.720067

**Authors:** Yosuke Teshirogi, Rina Mihara, Yuto Saito, Hyun-woo Rhee, Tohru Terada, Shin-ichi Tate, Kyota Yasuda

**Author notes:** Correspondence: Shin-ichi Tate, Kyota Yasuda. Equally contributed (Yosuke Teshirogi, Kyota Yasuda).

## Abstract

Stress granules (SGs) are dynamic, membrane-less assemblies that form in the cytoplasm in response to cellular stress. The ordered recruitment of proteins into SGs is fundamental to condensate composition and function, yet the molecular determinants of this ordered client recruitment remain incompletely understood. Using proximity photo-crosslinking proteomics, we identified heterogeneous nuclear ribonucleoprotein A2B1 (hnRNPA2B1) as a TIA1-proximal protein preferentially enriched in SGs under arsenite stress. Knockdown of hnRNPA2B1 preferentially delayed TIA1 enrichment in G3BP1-marked SGs at 20 min without affecting G3BP1 or the overall SG-positive cell fraction, and this phenotype showed directional rescue upon re-expression. In vitro droplet reconstitution assays with purified proteins revealed that hnRNPA2B1 and RNA cooperatively increased TIA1 incorporation capacity into G3BP1 condensates, an effect not attributable to changes in droplet size. Kinetic fitting identified hnRNPA2B1 + RNA as uniquely increasing the plateau amplitude of TIA1 recruitment (Cohen’s d = 1.62 versus RNA-alone condition). Coarse-grained simulations support an inside-out assembly model in which hnRNPA2B1 stabilizes the condensate core through homotypic interactions while RNA-bound TIA1 accumulates at the periphery. Together, these findings identify hnRNPA2B1 as a capacity-determining modulator of early TIA1 recruitment and provide a framework for understanding ordered protein assembly within stress granules.

## Introduction

Stress granules (SGs) are cytoplasmic, membrane-less organelles that assemble when cells are exposed to environmental stress ^1,2^. Their formation is initiated by the phase separation of RNA-binding proteins (RBPs), most notably G3BP1, which undergoes RNA-induced conformational changes that drive condensate nucleation ^3,4^. Subsequent recruitment of additional RBPs and mRNAs shapes the final composition of mature SGs, which is thought to be functionally relevant for the cellular stress response ^5,6^. Despite significant progress in understanding SG nucleation, the mechanisms that control ordered client recruitment during the early phase of SG assembly remain poorly characterized.

Recent studies have shifted current models of stress granule assembly toward a more RNA-centered view. G3BP1 has been shown to promote intermolecular RNA–RNA interactions during RNA condensation, functioning as an “RNA condenser” that stabilizes RNP granules ^7^. Furthermore, transcriptome-wide mRNP condensation can precede the appearance of microscopically visible stress granules ^8^, and a recent meta-analysis of SG transcriptome datasets has highlighted the complexity of the SG RNA world across conditions ^9^. Together, these findings support a hierarchical assembly model in which scaffold proteins, mRNPs, and client RBPs are incorporated in a temporally structured manner. Within this framework, factors that bias the early entry of specific RBPs into nascent stress granules remain incompletely defined.

TIA1 is a well-characterized SG component whose prion-related domain (PRD) mediates its aggregation and condensate incorporation ^10^. Although TIA1 can drive SG formation when overexpressed, it is not essential for G3BP1-positive SG formation under arsenite stress ^3^, suggesting that its recruitment into forming SGs may depend on auxiliary factors. Identifying such factors could reveal general principles of compositional control in biomolecular condensates.

To identify TIA1-associated proteins in the context of SG assembly, we applied proximity photo-crosslinking mass spectrometry using the Spotlight method ^11^. This strategy uses an azide-based Halo-ligand (VL1) to covalently crosslink proteins within approximately 1 nm of the bait protein upon UV irradiation, enabling detection of near-direct interaction partners with higher spatial resolution than conventional proximity labeling. Through this approach, we identified heterogeneous nuclear ribonucleoprotein A2B1 (hnRNPA2B1) as a TIA1-proximal candidate selectively enriched under stressed conditions.

hnRNPA2B1 is a multifunctional RBP involved in nuclear RNA processing, mRNA transport, stability, and m6A-dependent RNA metabolism ^12,13^. It localizes to SGs and has recently been shown to suppress arsenite-induced SG disassembly by maintaining the G3BP1–USP10/Caprin-1 interaction network ^14^. Additionally, hnRNPA2B1 has been identified as a linker connecting TIA1 and tau with m6A-modified RNA transcripts within condensates ^15^. Despite these connections, the role of hnRNPA2B1 in early TIA1 recruitment during SG assembly has not been directly examined.

Here, we report that knockdown of hnRNPA2B1 preferentially delays TIA1 enrichment in G3BP1-marked SGs at early time points without significantly affecting G3BP1. In vitro droplet assays confirm that hnRNPA2B1 promotes TIA1 incorporation into condensates cooperatively with RNA, acting as a capacity-determining factor rather than a rate accelerator. Coarse-grained simulations provide a structural basis for this RNA-mediated mechanism. Together, these findings identify hnRNPA2B1 as a modulator of early TIA1 recruitment and contribute to our understanding of the ordered assembly of biomolecular condensates.

## Results

### hnRNPA2B1 was identified as a TIA1-proximal protein enriched in stress granules

To identify proteins that interact with TIA1 in the context of SG formation, we employed proximity photo-crosslinking combined with biochemical SG core isolation and mass spectrometry. Halo-Flag-TIA1 expressed in HEK293 cells formed granular structures upon arsenite stress, and VL1 co-localized with these granules (Fig. S1A), confirming SG-dependent crosslinking. Using label-free quantification across three independent trials, we applied stringent filtering: proteins were retained only if detected in all three trials with UV+/UV− ratio >2-fold. After subtracting proteins common to stressed and non-stressed conditions, 14 candidate proteins selectively enriched under stress were identified (Table S2). STRING network analysis revealed hnRNPA2B1 as the only candidate with known physical interactions connecting it to multiple other candidates under stress (Fig. 1A, S1B). Time-course analysis showed hnRNPA2B1 was among the most persistently enriched proteins across the 20–60 min arsenite treatment window (Fig. 1B, C). These data identified hnRNPA2B1 as the top candidate for a TIA1-proximal protein in SGs.

**Figure 1.**
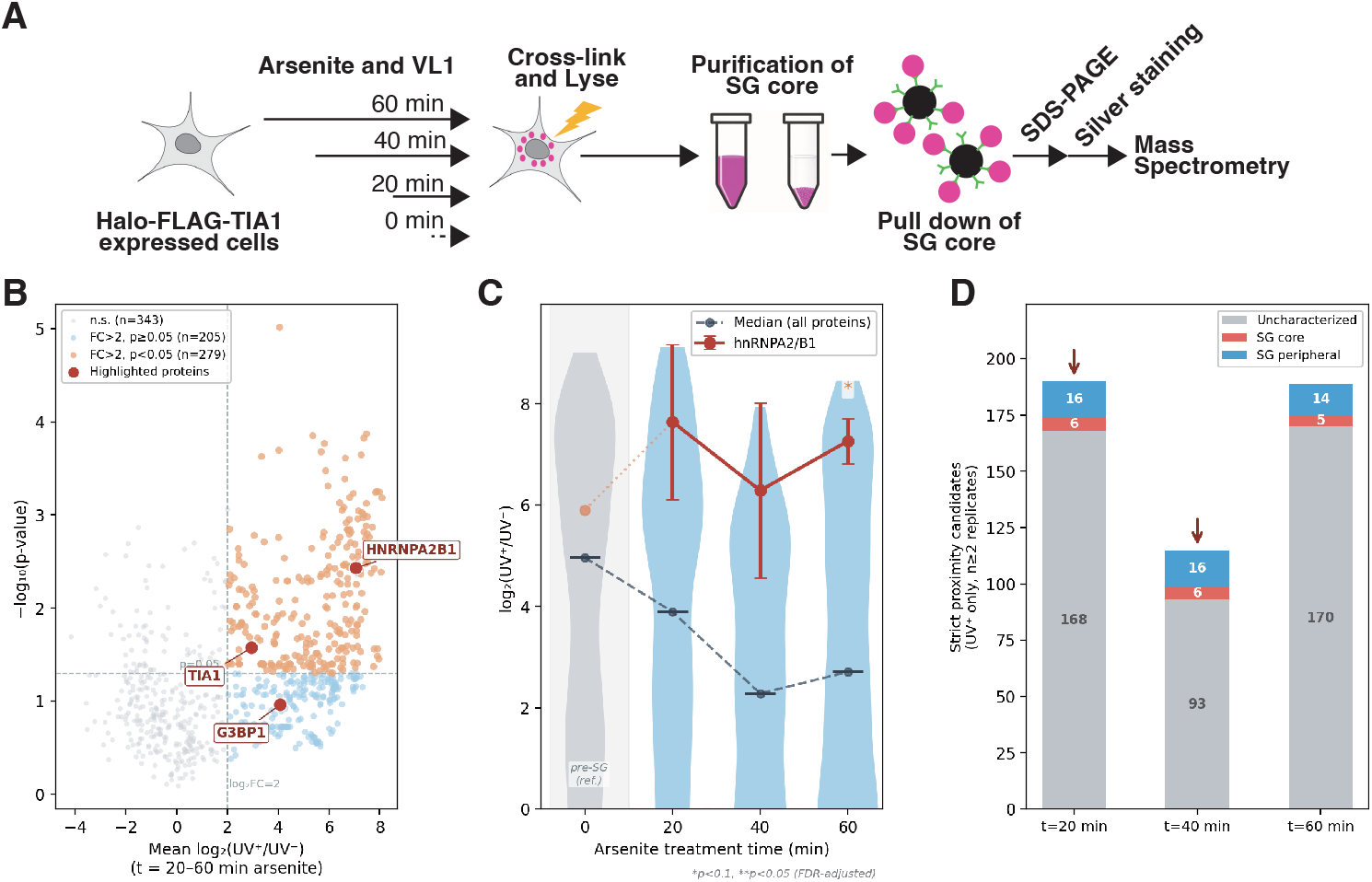
Identification of hnRNPA2B1 as a TIA1-proximal protein in stress granules by proximity photo-crosslinking mass spectrometry. (A) Schematic of the proximity photo-crosslinking and SG core mass spectrometry workflow. Halo-Flag-TIA1-expressing HEK293 cells were treated with VL1 and arsenite for the indicated times, subjected to UV-induced crosslinking, and processed for SG core isolation and mass spectrometry. (B) Volcano plot showing log2 fold-change (UV+/UV−) versus −log10(p-value) across all time points for detected proteins. Dashed lines indicate the log2FC = 2 and p = 0.05 thresholds. hnRNPA2B1, TIA1, and G3BP1 are highlighted. (C) Temporal enrichment profiles of hnRNPA2B1 and the median of all detected proteins across the arsenite time course. (D) Stacked bar charts showing the number of strict proximity candidates at each time point, categorized as SG core, SG peripheral, or uncharacterized.

### Knockdown of hnRNPA2B1 preferentially delays TIA1 recruitment into stress granules at early time points

To test whether hnRNPA2B1 influences TIA1 recruitment, we generated stable hnRNPA2B1 knockdown (A2B1-KD) HEK293 cells using lentiviral shRNA delivery. Knockdown efficiency was confirmed by western blot (Fig. S2A, B). Western blot analysis showed that TIA1 and G3BP1 protein levels were not significantly altered in KD cells (Fig. S2C, D), indicating that differences in SG composition reflect changes in recruitment rather than expression. hnRNPA2B1 localized to G3BP1-positive SGs under arsenite stress in control cells (Fig. 2A and Fig.S2E), consistent with its role as an SG component.

**Figure 2.**
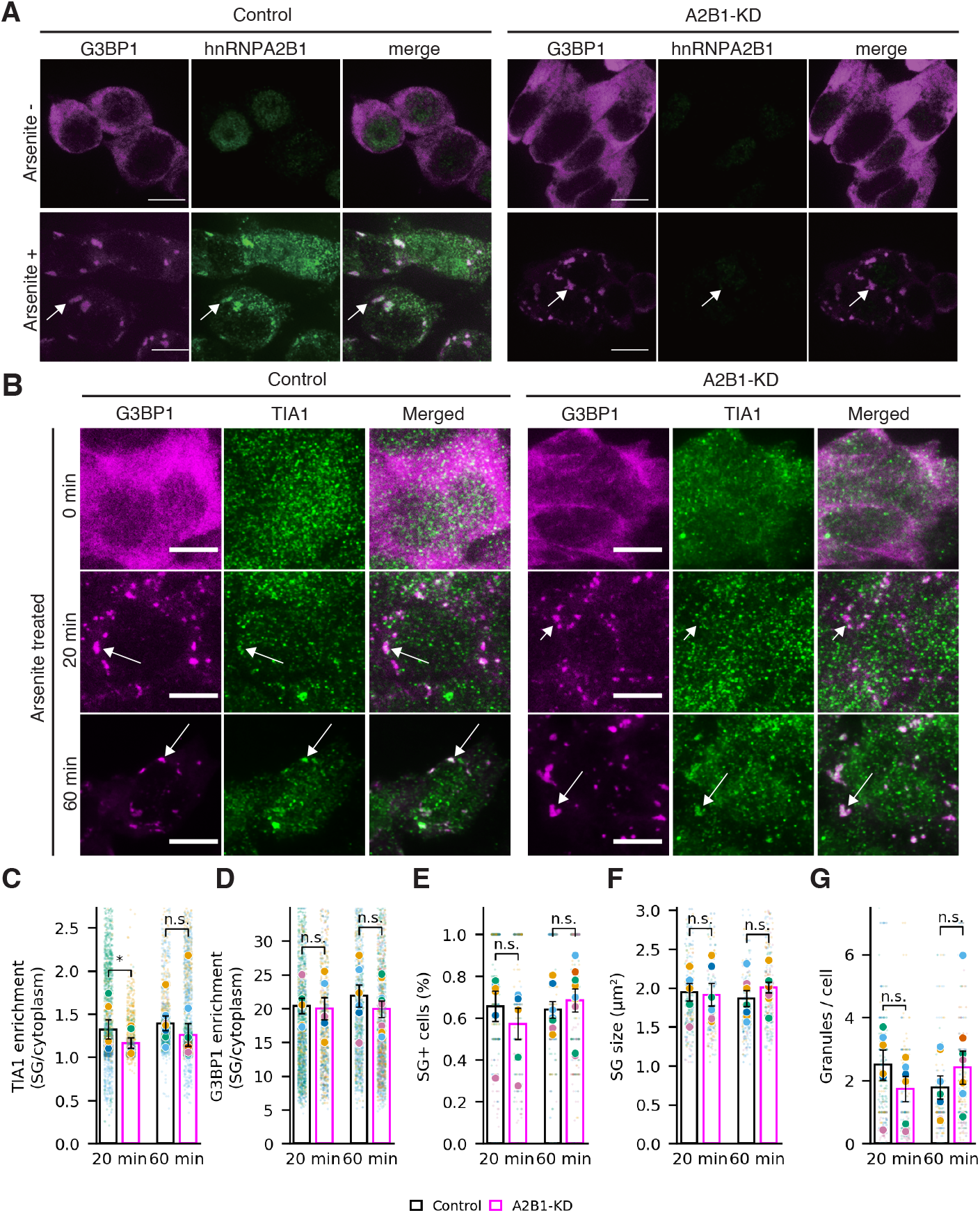
hnRNPA2B1 knockdown preferentially delays TIA1 recruitment into G3BP1-marked stress granules. (A) Representative immunofluorescence images of hnRNPA2B1 and G3BP1 in control and A2B1-KD cells with or without arsenite stress. (B) Representative time-course images of endogenous TIA1 and G3BP1 in control and A2B1-KD cells at 0, 20, and 60 min arsenite. (C) Super plots of TIA1 enrichment in G3BP1-marked SGs at 20 and 60 min. (D) As in (C), for G3BP1 enrichment. (E–G) Super plots showing SG-positive cell proportion, SG size, and granule number per cell. Small dots indicate individual granules or cells, large dots indicate replicate means, and bars indicate mean ± SEM. * p < 0.05 by paired t-test on replicate means.

To assess TIA1 and G3BP1 recruitment dynamics, control and A2B1-KD cells were subjected to arsenite stress for 20 or 60 min, fixed, and immunostained (Fig. 2B). TIA1 enrichment within G3BP1-positive SGs was significantly reduced in A2B1-KD cells at 20 min (mean difference = 0.171, 95% CI [0.042, 0.299], paired t-test p = 0.040, Cohen’s d = 1.06, n = 6 trials; Fig. 2C), with no significant difference at 60 min (p = 0.69, d = 0.17). G3BP1 enrichment was not significantly different at either time point (20 min: p = 0.71; 60 min: p = 0.26; Fig. 2D), indicating selectivity for TIA1. SG-positive cell fraction, SG size, and granule number per cell were not significantly different between conditions at either time point (Fig. 2E–G). These results indicate that hnRNPA2B1 promotes TIA1 entry into SGs specifically during early SG assembly.

### Re-expression of hnRNPA2B1 supports the specificity of the early TIA1 recruitment phenotype

To assess phenotype specificity, we introduced a codon-optimized, shRNA-resistant mKate-hnRNPA2B1 rescue construct into A2B1-KD cells and compared TIA1 enrichment at 20 and 60 min arsenite stress across control, KD, and rescue cells (Fig. S3). Control and KD cells expressed mKate alone; rescue cells expressed mKate-hnRNPA2B1. At 20 min, TIA1 enrichment was significantly lower in KD(mKate−) cells than in control(mKate−) cells (p = 0.008, d = 6.29, n = 3 trials). The rescue(mKate+) group showed a directional trend toward recovery relative to KD(mKate−) (d = −0.99), though this did not reach significance (p = 0.229), likely reflecting the modest effect of re-expressed hnRNPA2B1 that did not concentrate strongly in SGs. No clear group difference was detected at 60 min, consistent with the transient phenotype observed in the primary dataset. Although the re-expressed protein did not concentrate strongly within SGs, the directional recovery at 20 min supports the specificity of the knockdown phenotype and is consistent with a role for hnRNPA2B1 in early TIA1 recruitment.

To determine whether hnRNPA2B1 directly influences TIA1 incorporation into condensates— independent of the complex cellular environment—we reconstituted G3BP1 droplets in vitro using purified components and examined TIA1 recruitment kinetics under defined conditions.

### hnRNPA2B1 promotes TIA1 recruitment into condensates in vitro cooperatively with RNA

We next asked whether hnRNPA2B1 can directly influence TIA1 incorporation into G3BP1 condensates in vitro using a minimal reconstitution system. TIA1 fluorescence intensity within RFP-G3BP1-positive droplets was monitored over 15 min under four conditions: ±hnRNPA2-MBP and ±RNA (Fig. 3B, C). To quantify recruitment kinetics, TIA1 intensity time courses were fit to a single-component exponential saturation model, y = A(1 − exp(−kt)), where A represents the plateau amplitude (incorporation capacity) and k represents the rate constant (Supplementary Table S3).

**Figure 3.**
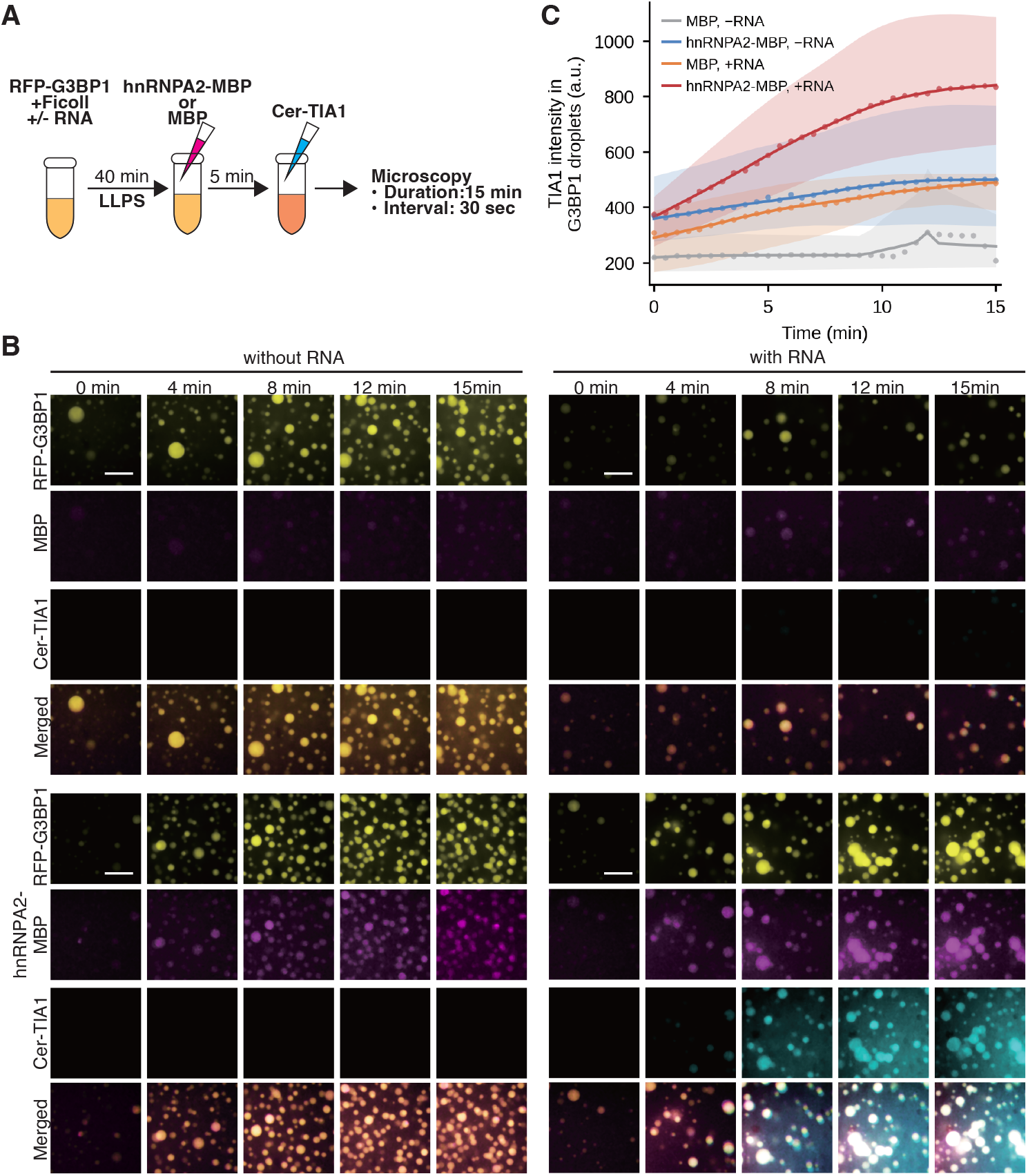
hnRNPA2 and RNA cooperatively enhance TIA1 incorporation into reconstituted G3BP1 condensates. (A) Schematic of the in vitro droplet reconstitution assay. RFP-G3BP1 (100 nM) was mixed with Ficoll (10%) ± RNA and incubated for 40 min to allow liquid-liquid phase separation. hnRNPA2-MBP or MBP (2 μM) was then added for 5 min, followed by addition of MBP-Cerulean-TIA1 (2 μM). Samples were immediately transferred to glass-bottom dishes for time-lapse imaging (30-second intervals, 15 min). (B) Representative time-lapse fluorescence images of reconstituted droplets under MBP (top) and hnRNPA2-MBP (bottom) conditions, without RNA (left) and with RNA (right). Channels: RFP-G3BP1 (yellow), hnRNPA2-MBP or MBP (magenta), MBP-Cerulean-TIA1 (cyan). Images are displayed at matched intensity ranges within each channel across all conditions to allow direct comparison. Scale bars, 10 μm. (C) Time course of MBP-Cerulean-TIA1 fluorescence intensity within RFP-G3BP1-positive droplets under four conditions. Data are LOESS-smoothed means with 95% bootstrap confidence intervals (5,000 resamples) of per-replicate means. Points indicate per-replicate means at each time point. n = 3 independent experiments. Kinetic parameters from exponential saturation fitting are reported in Supplementary Table S3.

The plateau amplitude A was greatest in the hnRNPA2-MBP + RNA condition (802 ± 252 a.u., mean ± SD, n = 3), compared with MBP + RNA (534 ± 137 a.u.), hnRNPA2-MBP alone (459 ± 205 a.u.), and MBP alone (242 ± 90 a.u.). Although the difference between the two RNA-containing conditions did not reach statistical significance (p = 0.181, n = 3), the effect size was large (Cohen’s d = 1.62), indicating that the modest sample size limits statistical power rather than reflecting an absent effect. The rate constant k was lower in the hnRNPA2-MBP + RNA condition (0.338 ± 0.020 min−1; t½ = 2.05 min) compared with conditions lacking hnRNPA2 or RNA, which showed faster but shallower saturation. This kinetic pattern—slower approach to a higher plateau—suggests that hnRNPA2 and RNA together increase the total capacity of the condensate to accommodate TIA1, rather than accelerating a capacity-limited process.

To address the possibility that larger droplets passively accumulate more TIA1 signal, we compared final droplet size between the two RNA-containing conditions. Droplet area did not differ between MBP + RNA and hnRNPA2-MBP + RNA (p = 0.736; Cohen’s d = 0.36, small effect), whereas TIA1 plateau amplitude showed a large effect size favoring the hnRNPA2-containing condition (Cohen’s d = 1.62). The dissociation between size and capacity effects indicates that enhanced TIA1 incorporation reflects a genuine increase in condensate recruitment capacity conferred by hnRNPA2 in an RNA-dependent manner.

The observation that hnRNPA2 increases TIA1 incorporation capacity while slowing saturation kinetics raises the question of whether these behaviors can be explained by the structural and dynamic properties of the proteins within the condensate. To investigate this, we performed coarse-grained molecular dynamics simulations.

### Coarse-grained simulations support an inside-out assembly model in which hnRNPA2B1 stabilizes the condensate core

To examine the physical basis of the recruitment phenotype, we performed coarse-grained simulations of multicomponent condensates containing G3BP1, TIA1, and hnRNPA2B1. Inclusion of hnRNPA2B1 increased the critical temperature (*T*_c_) of the simulated condensate (282.9 K) relative to the G3BP1-TIA1 system (271.1 K), indicating enhanced condensate stability (Fig. 4A). Spatial density analysis of the G3BP1-TIA1-hnRNPA2B1 system further showed that hnRNPA2B1 preferentially localized to the condensate core, while TIA1 was enriched toward the periphery and G3BP1 remained distributed throughout the condensate (Fig. 4B). Intermolecular contact analysis revealed that hnRNPA2B1 homotypic interactions were dominant, whereas G3BP1 maintained broad contacts with all molecular species (Fig. 4C). Together, these simulations support a model in which hnRNPA2B1 stabilizes the condensate core through self-association and promotes peripheral TIA1 recruitment.

**Figure 4.**
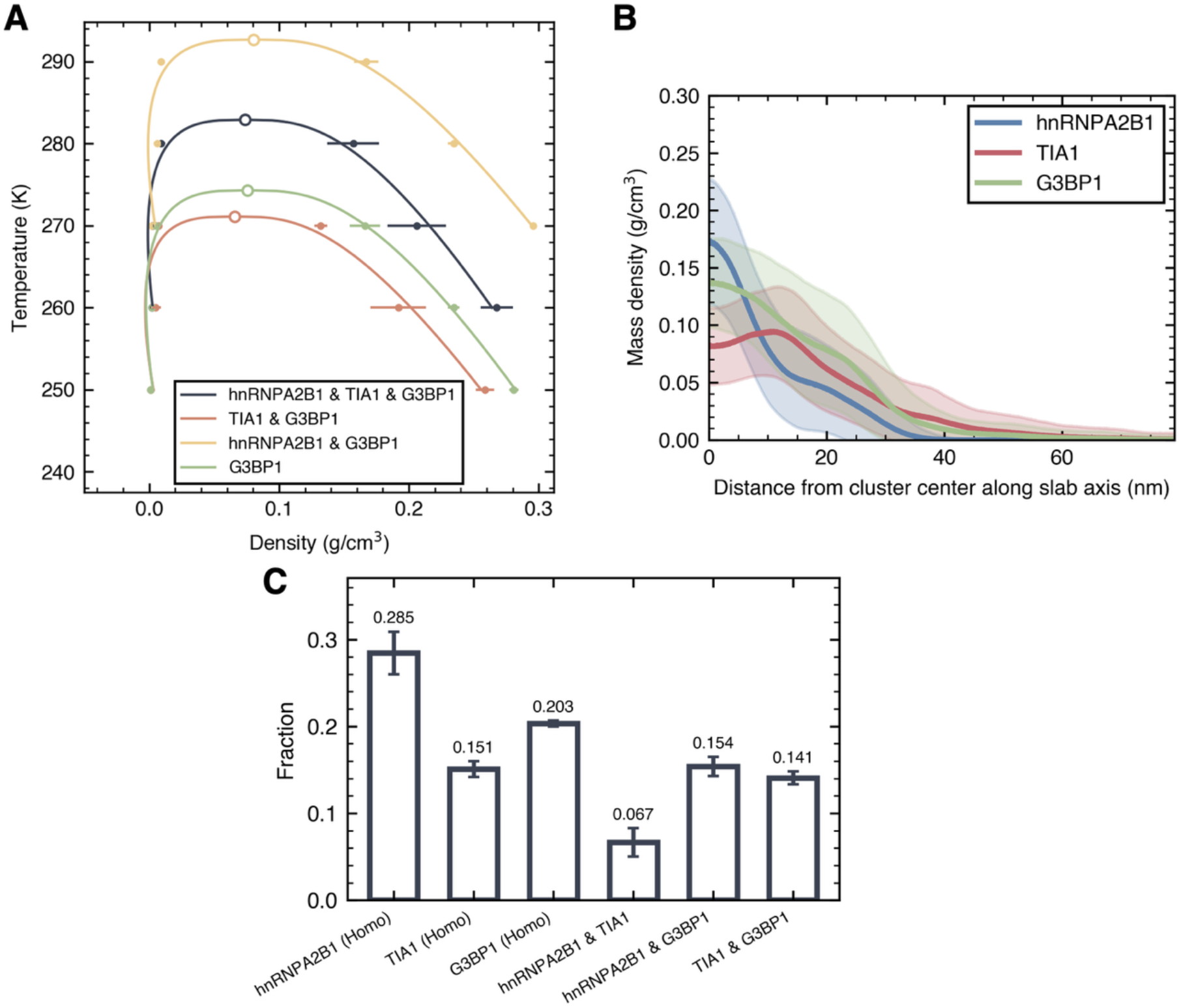
Coarse-grained simulations support an inside-out assembly model for hnRNPA2B1-dependent condensate organization. (A) Coexistence curves and corresponding critical temperatures for the simulated systems. Open circles indicate the critical points, and filled symbols with horizontal error bars show the mean ± SD of the densities of the high-density and low-density phases at the corresponding temperatures in three independent simulations. (B) Mass density profiles along the slab long axis, measured from the cluster center. Lines and shaded regions show the mean ± SD of three independent simulations. (C) Fractions of homo- and heterotypic intermolecular contacts relative to total contacts. Bars and error bars show the mean ± SD of three independent simulations.

## Discussion

Using proximity photo-crosslinking proteomics, cell-based perturbation, in vitro droplet reconstitution, and coarse-grained molecular dynamics simulations, we identified hnRNPA2B1 as a modulator of early TIA1 recruitment into stress granules. The most robust cellular phenotype was a selective reduction in TIA1 enrichment at 20 min of arsenite treatment after hnRNPA2B1 knockdown, with no comparable effect on G3BP1 and no clear difference at 60 min. Rescue analysis supported the specificity of this phenotype, and the in vitro and simulation datasets together provide a coherent framework for interpreting how hnRNPA2B1 influences ordered SG assembly.

A key point emerging from the cell-based analysis is that hnRNPA2B1 does not appear to be required for G3BP1-positive granule formation itself. Instead, it preferentially promotes efficient TIA1 entry during the early phase of SG assembly. This interpretation is consistent with the established view that G3BP1 serves as a core SG scaffold whereas TIA1 plays an important but modulatory role in SG formation ^3^. The temporal selectivity of the phenotype suggests that hnRNPA2B1 acts during the initial recruitment or stabilization of TIA1 at nascent granules rather than determining the eventual endpoint level of TIA1 accumulation.

The rescue experiment provides supportive, though incomplete, evidence for the specificity of the knockdown phenotype. Re-expression of shRNA-resistant mKate-hnRNPA2B1 produced a directional recovery of TIA1 enrichment at 20 min relative to KD cells (d = −0.99), suggesting that the phenotype reflects loss of hnRNPA2B1 function rather than off-target shRNA effects. However, the rescue was incomplete, likely because overexpressed hnRNPA2B1 did not localize efficiently to SGs—consistent with the known tendency of this predominantly nuclear protein to saturate nuclear import capacity upon overexpression ^14^. We therefore interpret the rescue data as consistent with, rather than definitive proof of, phenotypic specificity.

Our findings fit well with an emerging RNA-centered view of stress granule assembly. Recent work has shown that G3BP1 promotes intermolecular RNA–RNA interactions during RNA condensation and that widespread mRNP condensation can precede microscopically visible granule formation ^7,8^. In this view, an important unresolved question is not only how a granule core forms, but also how particular client RBPs enter this early assembly environment. Our data suggest that hnRNPA2B1 contributes at this level by promoting early TIA1 incorporation without broadly preventing G3BP1-positive granule formation. This interpretation is also consistent with recent evidence that G3BP-dependent stress granules reinforce the integrated stress response translation programme, supporting the idea that granule composition is functionally meaningful rather than merely structural ^6^.

The in vitro droplet assay indicates that hnRNPA2B1 and RNA cooperate to enhance TIA1 incorporation into G3BP1 condensates. Importantly, neither hnRNPA2B1 alone nor RNA alone reproduced the full effect observed when both were present, arguing for a cooperative mechanism rather than independent additive contributions. This result is consistent with the known RNA-binding properties of hnRNPA2B1 and with the broader idea that RBPs can shape condensate composition by organizing RNA-dependent interaction networks ^12,15^. Although the present data do not identify the relevant RNA species or prove direct molecular bridging, they support a model in which hnRNPA2B1 helps create an RNA-dependent context that favors TIA1 recruitment into forming SGs.

The kinetic analysis of in vitro TIA1 incorporation provides a more specific mechanistic interpretation. Fitting the time course to an exponential saturation model revealed that hnRNPA2B1 and RNA together selectively increase the plateau amplitude A—the maximum TIA1 incorporation capacity—without accelerating the rate constant k. Indeed, k was lower in the hnRNPA2B1 + RNA condition (t½ = 2.05 min) than in conditions lacking either component, indicating that TIA1 accumulates more slowly but reaches a substantially greater final level. We propose that hnRNPA2B1 acts primarily as a capacity-determining factor: it extends the TIA1-accessible volume or interaction surface within the condensate, rather than catalyzing rapid recruitment. This capacity model also provides a coherent interpretation of the cellular data. The significant reduction in TIA1 enrichment at 20 min after hnRNPA2B1 knockdown likely reflects a reduction in condensate capacity that is detectable when condensates have not yet reached saturation. At 60 min, compensatory pathways may suffice to populate TIA1 to a level that obscures the capacity difference, explaining the absence of a significant difference at the later time point. The identity of these compensatory mechanisms remains to be established. In this sense, hnRNPA2B1 may function less as a kinetic trigger of TIA1 entry and more as a determinant of how much TIA1 the early condensate can ultimately accommodate. Notably, final droplet size did not differ between conditions (p = 0.736; d = 0.36), ruling out passive size-dependent TIA1 accumulation as an alternative explanation.

A potential alternative interpretation of the in vitro data is that hnRNPA2B1 and RNA promote formation of larger droplets, which passively contain more TIA1 signal. This possibility is addressed directly by comparing droplet size and TIA1 capacity between RNA-containing conditions. Final droplet area did not differ between MBP + RNA and hnRNPA2-MBP + RNA (Cohen’s d = 0.36, small effect), whereas TIA1 plateau amplitude showed a large effect size favoring the hnRNPA2-containing condition (Cohen’s d = 1.62). The dissociation between these two metrics argues that the enhanced TIA1 incorporation reflects a change in condensate composition or interaction network rather than a trivial increase in condensate volume.

The simulation results refine this picture by suggesting an ‘inside-out’ organization principle (Figure 4). In this model, hnRNPA2B1 contributes to formation of a more stable condensate core through dominant homotypic interactions, whereas TIA1 preferentially accumulates in a more peripheral zone. This organization offers a plausible explanation for the selective, early, and transient phenotype observed in cells: perturbing hnRNPA2B1 would be expected to weaken the early molecular context that enables efficient TIA1 capture without necessarily preventing later-stage condensate maturation. The slower but higher-capacity kinetics observed in vitro (lower k, higher A) are consistent with a scenario in which hnRNPA2B1 first organizes the condensate core, creating a more favorable environment for TIA1 accumulation at the periphery over time. This localization of hnRNPA2B1 and TIA1 is consistent with the spatial density obtained from coarse-grained simulations.

Our findings are complementary to the recently reported role of hnRNPA2B1 in suppressing SG disassembly by maintaining the G3BP1-USP10-Caprin-1 interaction network ^14^. Together, these observations suggest that hnRNPA2B1 contributes to SG dynamics at multiple stages: first by promoting preferential early TIA1 entry during the initial phase of assembly, and later by helping preserve SG integrity during disassembly. This view places hnRNPA2B1 as a broader regulator of condensate composition and dynamics rather than a static structural component, consistent with emerging evidence that the balance between condensate assembly and clearance factors is fundamental to SG biology ^9,16^.

Several limitations of the present study should be acknowledged. The rescue experiment was complicated by incomplete SG localization of re-expressed mKate-hnRNPA2B1, likely attributable to saturation of nuclear import capacity at supraphysiological expression levels, and the proportion of SG-positive cells was lower in rescue cells than in controls (54% vs. 86% at 20 min), which may reflect a secondary effect of hnRNPA2B1 overexpression on SG dynamics and reduces the statistical power of the rescue comparison. The identity of the RNA species that mediates the cooperative effect of hnRNPA2B1 and RNA on TIA1 incorporation remains unknown, as do the compensatory mechanisms that normalize TIA1 enrichment at 60 min in knockdown cells. The sample sizes in the present study are modest (n = 6 trials for the primary knockdown analysis, n = 3 for rescue and in vitro experiments), and although effect sizes were reported to supplement p values, findings should be interpreted with appropriate caution pending independent replication. Finally, all cell-based experiments were performed in HEK293T cells under arsenite-induced stress; whether the identified role of hnRNPA2B1 generalizes to other cell types or stress conditions remains to be established.

In summary, this study identifies hnRNPA2B1 as a modulator of early TIA1 recruitment into stress granules and supports an RNA-dependent, capacity-based model for ordered SG assembly. The work does not yet define the exact interaction interface, the relevant RNA species, or the domain requirements underlying the recruitment effect. Those questions are best addressed in follow-up mechanistic studies. Nevertheless, the present data provide a coherent framework linking TIA1-proximal proteomics, selective cellular phenotypes, reconstitution, and simulation into a testable model of SG recruitment logic.

## Materials and Methods

### Plasmids and constructs

Halo-Flag-TIA1 was constructed in pcDNA3 using the human TIA1 coding sequence amplified from pF1KE0654 (KAZUSA DNA Research Institute) with EcoRI and XbaI sites. GFP-TIA1 was generated by cloning the same coding sequence into EGFP-C1 with SacI and BamHI sites. GFP-G3BP1 was constructed in EGFP-C1 using human G3BP1 amplified from cDNA prepared from HEK293 total RNA. The shRNA expression vector carried the shRNA cassette under the U6 promoter together with a puromycin-resistance gene (VectorBuilder). The hnRNPA2B1 target sequence was 5’-AGCTTCAGGTTATCGAAATA-3’, and the scramble control sequence was 5’-CCTAAGGTTAAGTCGCCCTCG-3’. Lentiviral packaging plasmids pMD2.G and psPAX2 were used for virus production. For rescue experiments, the human hnRNPA2 coding sequence was amplified from pJ4M_hnRNPA2_FL_WT, a gift from Nicolas Fawzi (Addgene plasmid #139109; http://n2t.net/addgene:139109; RRID:Addgene_139109) ^17^, codon-optimized with synonymous substitutions across the shRNA target region, and cloned into a mammalian expression vector with an N-terminal mKate2 tag. For recombinant protein production, mRFP1-G3BP1 was constructed in pET28a by inserting mRFP1 and human G3BP1 coding sequences. MBP-Cerulean-TIA1 was custom-synthesized in a bacterial expression vector (VectorBuilder). MBP alone was expressed from pET28a into which the MBP coding sequence was cloned from pMAL-c6T (N0378S, New England Biolabs). hnRNPA2-MBP was expressed from pJ4M_hnRNPA2_FL_WT.

### Recombinant protein expression and purification

All recombinant proteins were expressed in E. coli BL21(DE3). Cultures were grown at 37°C in LB medium until OD600 reached 0.6–0.8, induced with 1 mM IPTG for 4 h at 37°C, and further cultured for 16–20 h at 16°C.

MBP-Cerulean-TIA1 was grown in carbenicillin (50 μg/ml). Cells were resuspended in Column Buffer (20 mM Tris-HCl pH 7.4, 200 mM NaCl, 1 mM DTT, 10% glycerol), lysed by sonication, and centrifuged. The clarified lysate was loaded onto amylose resin, washed with Column Buffer, and eluted with Column Buffer supplemented with 10 mM maltose. Protein was dialyzed into HEPES Buffer (20 mM HEPES-NaOH pH 7.4, 200 mM NaCl) at 4°C. ε = 167,560 M−1cm−1, MW = 111.0 kDa.

RFP-G3BP1 and hnRNPA2-MBP were grown in kanamycin (20 μg/ml). Cells were resuspended in Buffer A1 (50 mM Tris-HCl pH 9.0, 200 mM NaCl, 4 M urea, 10 mM imidazole), lysed by sonication, and centrifuged. The clarified lysate was loaded onto Ni-NTA resin, washed sequentially with Buffer A2 (2 M urea, 10 mM imidazole), Buffer B (4 M NaCl, 2 M urea, 10 mM imidazole), and Buffer C (50 mM NaCl, 2 M urea, 10 mM imidazole; all in 50 mM Tris-HCl pH 9.0), then eluted with 500 mM imidazole. Proteins were dialyzed against 50 mM Tris-HCl pH 9.0, concentrated to 5 ml, purified by size-exclusion chromatography in 50 mM Tris-HCl pH 7.4, and dialyzed into HEPES Buffer at 4°C. RFP-G3BP1: ε = 49,640 M−1cm−1, MW = 80.2 kDa. hnRNPA2-MBP: ε = 101,130 M−1cm−1, MW = 80.1 kDa.

RFP-G3BP1, hnRNPA2-MBP, and MBP were grown in kanamycin (20 μg/ml). Cells were resuspended in Buffer A1 (50 mM Tris-HCl pH 9.0, 200 mM NaCl, 4 M urea, 10 mM imidazole), lysed by sonication, and centrifuged. The clarified lysate was loaded onto Ni-NTA resin, washed sequentially with Buffer A2 (2 M urea, 10 mM imidazole), Buffer B (4 M NaCl, 2 M urea, 10 mM imidazole), and Buffer C (50 mM NaCl, 2 M urea, 10 mM imidazole; all in 50 mM Tris-HCl pH 9.0), then eluted with 500 mM imidazole. Proteins were dialyzed against 50 mM Tris-HCl pH 9.0, concentrated to 5 ml, purified by size-exclusion chromatography in 50 mM Tris-HCl pH 7.4, and dialyzed into HEPES Buffer at 4°C. Molar extinction coefficients and molecular weights were: RFP-G3BP1, ε = 49,640 M−1cm−1, MW = 80.2 kDa; hnRNPA2-MBP, ε = 101,130 M−1cm−1, MW = 80.1 kDa; MBP, ε = 67,965 M−1cm−1, MW = 48.4 kDa.

### Cell culture and transfection

HEK293T cells (RIKEN Cell Bank, RCB2202) were maintained in DMEM supplemented with 10% fetal bovine serum, penicillin, and streptomycin at 37°C in 5% CO2. Stable knockdown cells (A2B1-KD) were generated by lentiviral transduction and selected with 2 μg/ml puromycin. For transient transfection, plasmids were introduced using PEI MAX in Opti-MEM, and cells were analyzed 12 h after transfection unless otherwise indicated. Oxidative stress was induced by treatment with 0.5 mM sodium arsenite for the times indicated in each experiment.

### Proximity photo-crosslinking and SG core isolation

Spotlight proximity photo-crosslinking was performed essentially as described previously ^11,18^. Halo-Flag-TIA1-expressing HEK293T cells were incubated with VL1 (10 μM, 1 h) at 37°C with or without 500 μM sodium arsenite, rinsed twice with PBS, and subjected to UV irradiation at 20,000 μJ/cm^2^ for 10 min (UVP CL-1000, Analytik Jena). Cells were pelleted by centrifugation (1,500 × g, 3 min), snap-frozen in liquid nitrogen, and stored at −80°C until use.

SG core isolation followed a previously reported protocol ^19^. Cell pellets were resuspended in SG lysis buffer (50 mM Tris-HCl pH 7.4, 100 mM potassium acetate, 2 mM magnesium acetate, 0.5 mM DTT, 50 μg/ml heparin, 0.5% NP-40, 1 U/μl RNase inhibitor, and protease inhibitor) and lysed by passage through a 25-gauge needle seven times on ice. Lysates were centrifuged at 1,000 × g for 5 min to remove debris, and the supernatant was centrifuged at 18,000 × g for 20 min. The pellet was washed and centrifuged at 900 × g for 2 min to obtain the SG core fraction. The supernatant was immunoprecipitated with anti-FLAG antibody-conjugated Dynabeads (Fujifilm Wako, cat# 017-25151) for 1 h at 4°C, washed sequentially with Buffer 1 (20 mM Tris-HCl pH 8.0, 500 mM NaCl), Buffer 2 (20 mM Tris-HCl pH 8.0, 200 mM NaCl), and Buffer 3 (10 mM Tris-HCl pH 8.0, 250 mM LiCl, 1 mM EDTA), and eluted with 2× Laemmli sample buffer at 98°C for 5 min. Samples lacking UV irradiation were processed in parallel as negative controls.

### Mass spectrometry and proteomic analysis

Eluted proteins were reduced, alkylated, and digested with trypsin by in-gel digestion ^20^. Peptides were analyzed by nanoLC-MS/MS using an LTQ Orbitrap XL mass spectrometer (Thermo Fisher Scientific) coupled to an Ultimate 3000 RSLCnano system (Thermo Fisher Scientific). Raw files were processed and annotated by peptide mass fingerprinting using CHIMERYS in Proteome Discoverer version 3.0 (Thermo Fisher Scientific) against the UniProt human database. Trypsin specificity was set with a maximum of two missed cleavages; precursor mass tolerance was 10 ppm and fragment mass tolerance was 0.8 Da. Missing values in the UV− channel were imputed using the sample minimum (SampMin) method ^21^, consistent with a missing-not-at-random assumption for proteins absent from the non-crosslinked control. Candidate proteins were retained only when detected in three to four independent biological replicates with a UV+/UV− ratio of at least twofold at one or more time points, with n ≥ 2 replicates per time point. Statistical significance was assessed by one-sample t-test against zero, with Benjamini–Hochberg false discovery rate correction applied across all detected proteins ^22^. Proteins detected under both stressed and non-stressed conditions were removed from the final candidate list. The complete list of detected proteins is provided in Supplementary Table S1. The 14 final candidates are listed in Supplementary Table S2

### Immunostaining and western blotting

Primary antibodies included anti-hnRNPA2B1 (DF6296, Affinity Biosciences), anti-TIA1 (HPA056961-100UL, Sigma-Aldrich), anti-G3BP1 (ab56574, Abcam), anti-GAPDH (10494-1-AP, Proteintech). For western blotting, cells were lysed in RIPA buffer containing protease inhibitors, and lysates were quantified by BCA assay before SDS-PAGE and transfer to PVDF membranes. TIA1 was detected as a doublet at approximately 40–45 kDa corresponding to the two known isoforms ^23^; the upper doublet was used for quantification, whereas the lower approximately 37 kDa band was treated as non-specific. For immunofluorescence, cells were fixed in 4% formaldehyde, permeabilized in 0.2% Triton X-100, and blocked in 5% FBS in PBS. Samples were mounted in Fluoromount-G with DAPI. Uncropped western blot membrane images are provided in Figure S4.

### Fluorescence microscopy

Images were acquired on an Olympus IX83 inverted microscope equipped with a Yokogawa CSU-W1 spinning-disk confocal unit. Fixed samples were imaged using a 100× oil-immersion objective (NA 1.4). Excitation wavelengths were 405 nm for DAPI, 488 nm for GFP/CoraLite488/FITC/VL1, and 637 nm for Alexa Fluor 647. Z-stacks were collected at 0.2 μm intervals (40 optical sections for cells, 20 optical sections for *in vitro* droplet) and converted to maximum-intensity projections for analysis. For in vitro droplet imaging, samples were transferred to glass-bottom dishes and imaged using a 60× oil-immersion objective. Cerulean was excited at 445 nm, RFP at 561 nm, and Alexa Fluor 633 at 637 nm. Time-lapse images were acquired at 30-second intervals for 15 minutes (31 time points), with z-stacks of 20 optical sections converted to maximum-intensity projections for quantification.

### Image analysis and statistics

Stress granules were detected from the G3BP1 channel using a custom Python pipeline based on scikit-image. Images were processed by white top-hat filtering, Gaussian smoothing, and Otsu thresholding (multiplier = 1.68). Candidate regions were filtered by area, solidity, and eccentricity. For each SG, mean fluorescence intensity was measured within the SG mask and in a surrounding ring region, and enrichment was calculated as the SG/cytoplasm ratio. Cell masks were approximated by expanding DAPI-defined nuclear masks by 130 pixels. In rescue experiments, mKate-positive cells were identified using Otsu thresholding of mean cellular mKate intensity. Quantitative imaging data were analyzed using a super-plot framework in which biological replicate means were used as the unit of statistical testing, thereby avoiding pseudoreplication ^24^.

Paired t-tests were applied to replicate means, 95% confidence intervals were calculated from paired differences, and effect sizes were reported as Cohen’s d. Values exceeding the group mean ± 3 SD were excluded prior to analysis.

For in vitro droplet quantification, mean fluorescence intensity of MBP-Cerulean-TIA1 within RFP-G3BP1-positive droplets was measured at each time point. Droplets were segmented from the RFP channel by Otsu thresholding, and TIA1 signal within droplet masks was quantified across all 31 time points. For kinetic analysis, per-replicate mean intensities were fit to a single-component exponential saturation model, y = A(1 − exp(−kt)), using nonlinear least-squares optimization (scipy.optimize.curve_fit). The plateau amplitude A and rate constant k were extracted per replicate, and t½ was calculated as ln(2)/k. Time-course data in Fig. 3C are presented as LOESS-smoothed means (span = 0.35) with 95% bootstrap confidence intervals (5,000 resamples, percentile method) of per-replicate means. Statistical comparisons of kinetic parameters and final droplet size between conditions were performed by unpaired t-test. Effect sizes were reported as Cohen’s d to supplement p values given the limited sample size (n = 3 per group).

### In vitro droplet reconstitution assay

Recombinant condensates were assembled by mixing RFP-G3BP1 (100 nM final), RNA (100 ng/μL), and Ficoll (10% w/v) in HEPES Buffer (20 mM HEPES-NaOH pH 7.4, 200 mM NaCl) to 18 μL total volume, and incubating for 40 min to allow droplet formation. Alexa Fluor 633 C5-maleimide (A20342, Thermo Fisher Scientific)-labeled hnRNPA2-MBP or MBP (2 μM final, 1 μL) was then added and incubated for 5 min, followed by addition of MBP-Cerulean-TIA1 (2 μM final, 1 μL). Four conditions were compared: MBP, −RNA; hnRNPA2-MBP, −RNA; MBP, +RNA; and hnRNPA2-MBP, +RNA. Samples were immediately transferred to glass-bottom dishes for time-lapse imaging. TIA1 incorporation into G3BP1-positive droplets was quantified at each time point. Data represent three independent experiments.

### Coarse-grained simulations

Protein structural models (UniProtKB: P22626, P31483, Q13283) were obtained from AlphaFold2 DB^25^ and partitioned into folded and disordered regions. This classification was performed using a threshold of pLDDT = 70, based on previous research^26^. Residue pairs within folded domains with Cα-Cα distances of less than 10 Å were restrained using elastic network to maintain tertiary structure. This distance cutoff was determined to ensure G3BP1 NTF2-domain dimerization. The Mpipi-recharged residue-level potential^27^ was employed to model protein interactions. Each system contained 50 molecules per species for three-component systems and 64 molecules per species for two-component systems, arranged in a slab geometry. The Debye length was set to 7.93 Å corresponding to a salt concentration of 150 mM. Simulations were performed using a modified version of OpenMpipi^28^. Following the original OpenMpipi protocol, the initial condensed phase was initially prepared by pulling all particles gently toward the center of the slab box for 20 ns, after which production simulations were carried out with snapshots recorded every 500 ps. Simulations were conducted using Langevin dynamics with a friction coefficient of 0.01ps^-1^. For each condition, three independent 2 μs simulations were performed at temperatures ranging from 250 K to 310 K in 10 K intervals to estimate critical temperatures. The critical density and temperature (ρ_c_, *T*_c_) were estimated via fitting the coexistence data to the following curves:

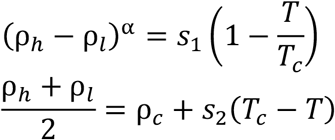

where ρ_*h*_ and ρ_*l*_ refer to the densities of the high-density phase and the low-density phase, respectively. *s*_1_ and *s*_2_ are fitting parameters, and α = 3.06. These settings were followed the original Mpipi-recharged model study^27^. Spatial density profiles were calculated along the slab long axis after centering the cluster. The fractions of molecular contact frequencies were calculated by summing residue-level contacts identified using a 10 Å cutoff and normalizing by the total number of contacts. Density and contact analysis were performed using trajectories at 270 K (approximately 0.95 *T*_c_). All coarse-grained simulations were carried out using OpenMM 8.2^29^ and analyzed with in-house code.

## Acknowledgements

The proteomics work was conducted with the facilities of the Natural Science Center for Basic Research and Development (N-BARD) at Hiroshima University [NBARD-00174]. We thank Dr. Nobuo Yamaguchi and Dr. Tomoko Amimoto of N-BARD at Hiroshima University for their contributions to the in-gel digestion and support for data acquisition by mass spectrometric analysis used in this work.

## Author contributions

S-iT and KY were involved in conceptualization; RM performed the mass spectrometry; KY and RM performed proteomic data analysis, KY performed cellular experiments, YS performed experiments with synthesized protein, KY performed all image analysis, YT performed simulation and analysis, H-WR and KY were involved in resources; KY wrote the original draft; KY, YT and TT were involved in writing – reviewing and editing; KY and ST were involved in project administration.

## Figure Legends

**Figure S1.**
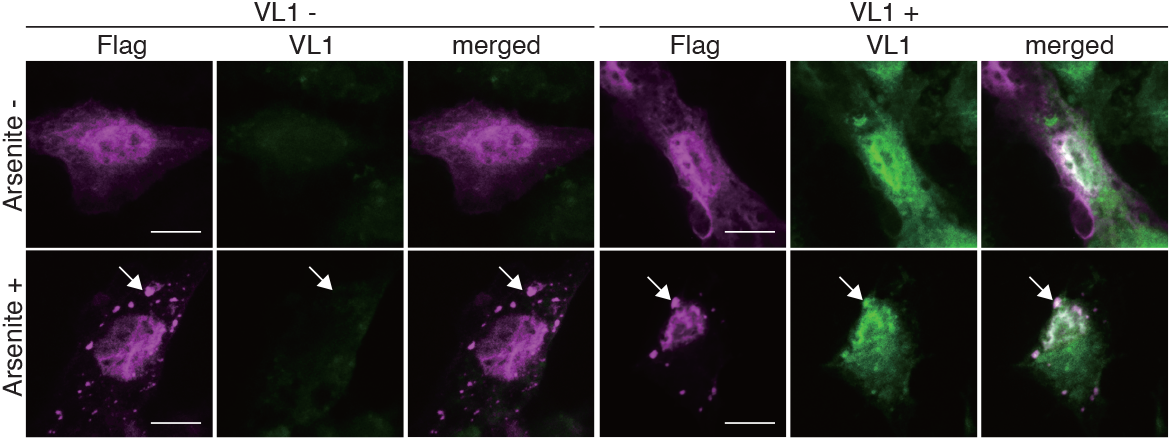
Validation of proximity photo-crosslinking in stress granules. Representative immunofluorescence images of Halo-Flag-TIA1 (anti-Flag, magenta) and VL1 autofluorescence (green) in HEK293 cells ± arsenite, ± VL1. Arrows indicate granular structures. VL1 co-localizes with Halo-Flag-TIA1 granules only in the arsenite + VL1 condition. Scale bar, 10 μm.

**Figure S2.**
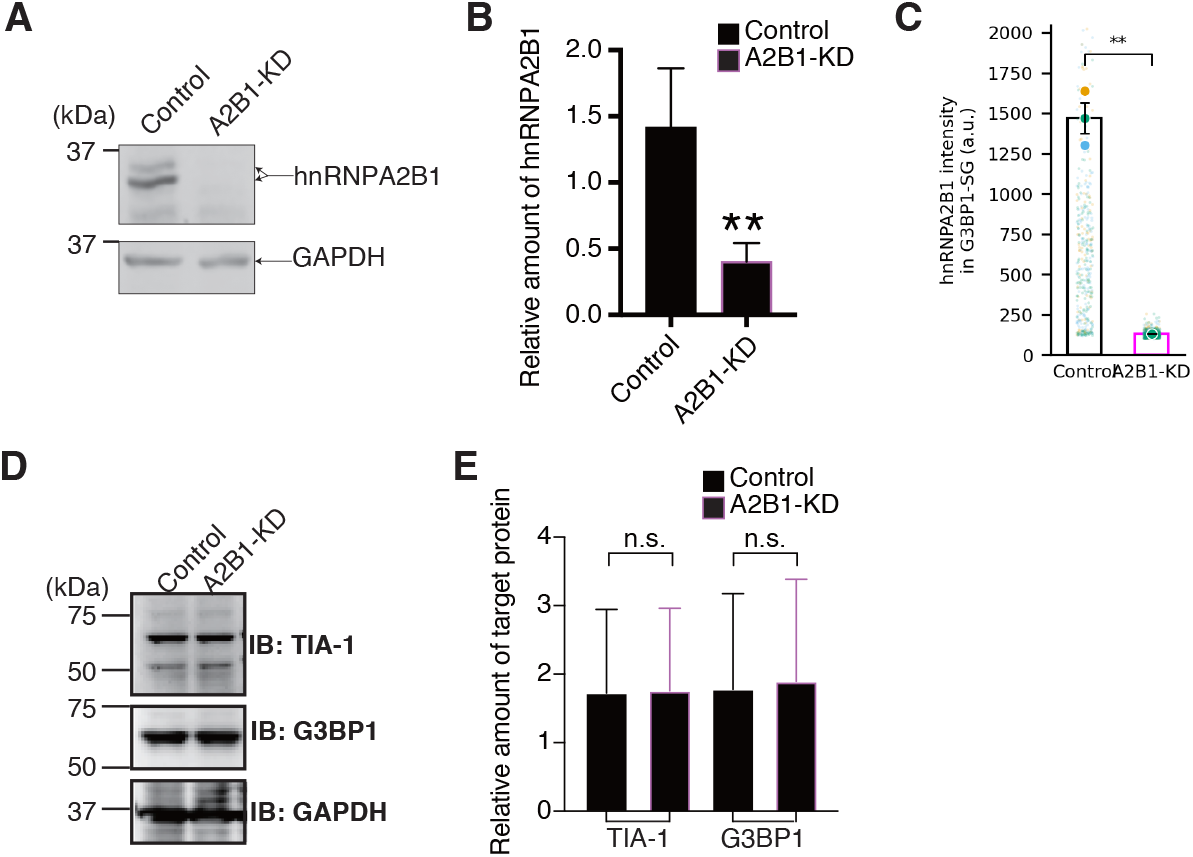
Validation of hnRNPA2B1 knockdown and basal expression levels of TIA1 and G3BP1. (A, B) Western blot and quantification of hnRNPA2B1 in control and A2B1-KD cells, normalized to GAPDH. ** p = 0.0034 by paired t-test. Mean ± SEM, n = 3. (C, D) Western blot and quantification of TIA1 and G3BP1. TIA1 was detected as a doublet at ∼40–45 kDa corresponding to the two known isoforms ^23^; the upper doublet was used for quantification; the lower ∼37 kDa band is non-specific. n.s. for both by paired t-test. Mean ± SEM, n = 3. (E) Super plot of hnRNPA2B1 fluorescence intensity within G3BP1-positive SGs in control cells under arsenite stress. ** p = 0.0052 by paired t-test. n = 3 trials.

**Figure S3.**
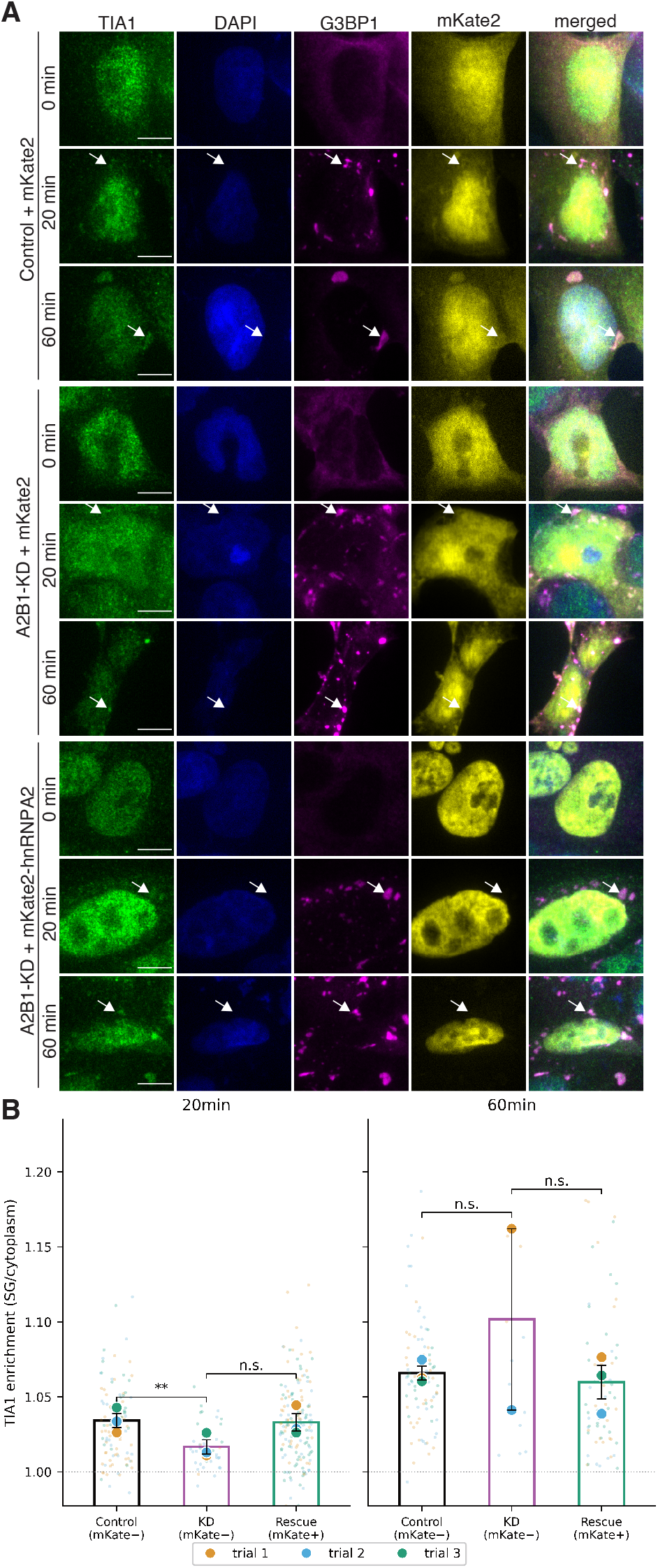
Re-expression of hnRNPA2B1 partially restores early TIA1 recruitment in knockdown cells. Super plots of TIA1 enrichment in G3BP1-marked SGs at 20 and 60 min arsenite in control (mKate−), A2B1-KD (mKate−), and rescue (mKate-hnRNPA2B1+) cells. At 20 min, TIA1 enrichment was significantly lower in KD versus control (p = 0.008, Cohen’s d = 6.29). The rescue group showed directional recovery (d = −0.99) without reaching significance (p = 0.229, n = 3 trials). No significant difference was detected at 60 min. One trial was excluded based on predefined rescue-expression QC criteria.

**Figure S4.**
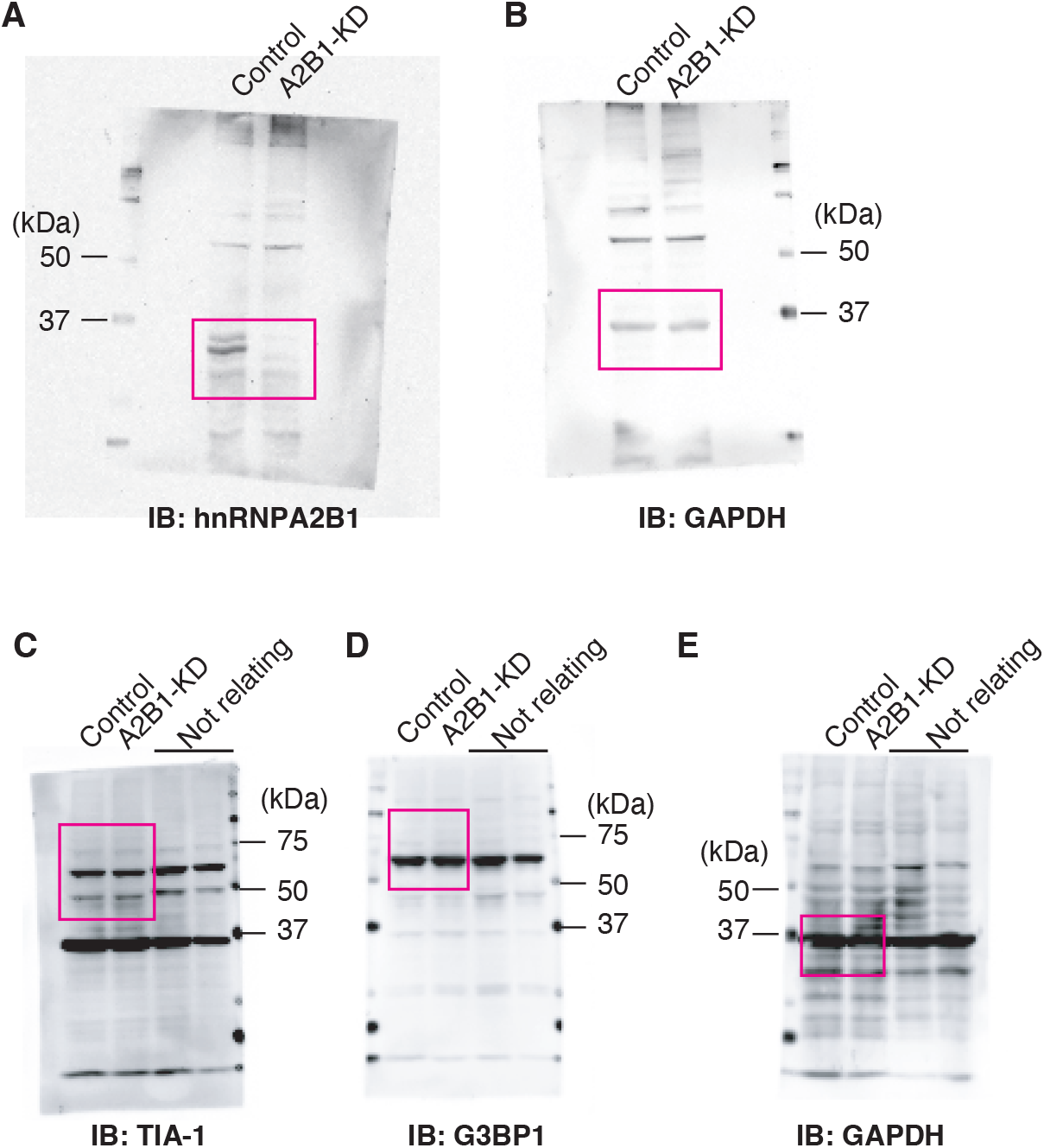
Uncropped western blot membranes. (A) Uncropped membrane for IB: hnRNPA2B1 (corresponding to Fig. S2A). (B) Uncropped membrane for IB: GAPDH (loading control for A). (C) Uncropped membrane for IB: TIA-1 (corresponding to Fig. S2C). (D) Uncropped membrane for IB: G3BP1 (corresponding to Fig. S2C). (E) Uncropped membrane for IB: GAPDH (loading control for C). Magenta boxes indicate the regions shown in the corresponding cropped panels. “Not relating” lanes contain samples unrelated to this study.

**Supplementary Table S1. Complete list of proteins detected by mass spectrometry**.

All 676 proteins identified across stressed and non-stressed conditions with their label-free quantification values.

**Supplementary Table S2. Candidate TIA1-proximal proteins identified by proximity photo-crosslinking mass spectrometry under stress**.

Proteins detected with UV+/UV− ratio ≥ 2-fold in ≥ 2 of 3–4 independent replicates under arsenite stress conditions, after removal of proteins common to stressed and non-stressed conditions. The 14 remaining candidates are listed with their average relative quantities.

**Supplementary Table S3. Kinetic parameters of TIA1 incorporation into reconstituted G3BP1 condensates in vitro**.

(A) Parameters from exponential saturation fitting [y = A(1 − exp(−kt))]. A, plateau amplitude; k, rate constant; t½, half-time. Mean ± SD, n = 3. (B) Pairwise statistical comparisons by unpaired t-test with Cohen’s d effect sizes.

